# Prevalence of the fungal pathogen *Batrachochytrium dendrobatidis* in amphibians of Costa Rica predated first-known epizootic

**DOI:** 10.1101/482968

**Authors:** Marina E. De León, Héctor Zumbado-Ulate, Adrián García-Rodríguez, Gilbert Alvarado, Hasan Sulaeman, Federico Bolaños, Vance T. Vredenburg

**Affiliations:** Department of Microbiology and Molecular genetics, University of California, Davis, USA; Department of Biological Sciences, Purdue University, West Lafayette, IN, USA; Escuela de Biología, Universidad de Costa Rica, San Pedro, Costa Rica; Departamento de Ecologia, Universidade Federal do Rio Grande do Norte, Natal - RN, Brazil; Faculdade de Medicina Veterinária e Zootecnia, Universidade de São Paulo, São Paulo, Brazil; Department of Biology, San Francisco State University, San Francisco, California, USA; Museum of Vertebrate Zoology, University of California Berkeley, Berkeley, California, USA

**Author notes:** These authors contributed equally to this work.

## Abstract

Emerging infectious diseases are a growing threat to biodiversity worldwide. Outbreaks of the infectious disease chytridiomycosis, caused by the fungal pathogen *Batrachochytrium dendrobatidis* (*Bd*), have caused the decline and extinction of numerous amphibian species. In Costa Rica, a major decline event occurred in 1987, more than two decades before this pathogen was discovered. The loss of many species in Costa Rica is assumed to be due to *Bd*-epizootics, but there are few studies that provide data from amphibians in the time leading up to the proposed epizootics. In this study, we provide new data on *Bd* infection rates of amphibians collected throughout Costa Rica, in the decades prior to the epizootics. We used a quantitative PCR assay to test for *Bd* infection in 1016 specimens collected throughout Costa Rica. We found *Bd*-infected hosts collected as early as 1964, and a infection prevalence average per decade of just 4%. The infection prevalence remained relatively low and geographically constrained until the 1980s when epizootics are hypothesized to have occurred. After that time, infection prevalence increased three-fold and *Bd*-infected hosts we collected throughout the entire country. Our results, suggest that *Bd* may either have invaded Costa Rica earlier than previously known, and spread more slowly than previously reported, or that an endemic lineage of the pathogen may exists. To help visualize areas where future studies should take place, we provide a *Bd* habitat suitability model trained with local data. Studies that provide information on genetic lineages of *Bd* are needed to determine whether an endemic lineage of *Bd* or the Global Panzootic Lineage (identified from mass die off sites globally) was present in Costa Rica and responsible for the epizootics that caused amphibian communities to collapse.

## Introduction

Amphibians are experiencing a global extinction event [1,2]. Though many factors contribute to population declines, the emergence of the fungal pathogen *Batrachochytrium dendrobatidis* (*Bd*) is one of the most important [3]. The disease chytridiomycosis, caused by the fungal pathogen *Batrachochytrium dendrobatidis* (hereafter *Bd*), was first described in 1999 and has since been found all over the world [3–5]. Interestingly, *Bd* is composed of many genetic lineages that vary in virulence and affect host species differently. The panzootic disease is attributed to *Bd*-GPL, a Global Panzootic Lineage of *Bd* associated with *Bd* epizootics and host population collapse [6]. Other lineages of *Bd* have been shown to be less virulent and have been identified in areas lacking epizootics [7]. *Bd* infects the skin of the amphibian and causes hyperkeratosis, the thickening of skin which disrupts the amphibian’s osmotic balance; leading to death by cardiac arrest in highly infected individuals [8,9].

The dynamics of *Bd* and its hosts, including pathogen invasion and the host-pathogen interactions that follow, are still not fully understood. For example, in some areas (e.g. South Korea, Brazil, and South Africa), *Bd* appears to be in an enzootic state with amphibian hosts [10–12], while in others (eg. western North America [13], Central America and South America) there are repeated examples of epizootics and die offs of hosts. In these areas, *Bd*-GPL is associated with epizootics [14]. South Korea was recently proposed as a region of high *Bd* genetic diversity, with one of the *Bd* lineages identified as exhibiting genetic hallmarks that may be the source of the panzootic *Bd* (*Bd*-GPL) that emerged in the 20^th^ century [15]. Many of the reported declines attributed to *Bd*-GPL in the new world occurred decades before *Bd* was described, thus, retrospective studies can help create a timeline for *Bd* emergence and spread.

Causes of amphibian declines in Costa Rica, where some of the earliest reported declines of amphibians occurred, have been debated in the literature [16,17]. Some studies proposed *Bd* epizootics occurred when environmental factors weakened host immune systems making them more susceptible to endemic *Bd* [18]. Other studies refute this and show that an invasive *Bd* pathogen caused the epizootics [19,20]. Costa Rica had one of the earliest amphibian declines (1980s and 1990s) that was later associated with *Bd* epizootics [21–23]. These declines mostly affected stream-dwelling species at elevations between 1000 and 2500 meters and include sites such as Monteverde, where the amphibian community collapsed a decade before *Bd* was described [24]. At this site, around the year 1987, half of all amphibian species, along with the Costa Rican golden toad (*Incilius periglenes*), disappeared [22].

Like many other areas experiencing *Bd* epizootics, anuran (frogs and toads) species in Costa Rica experienced differential susceptibility to *Bd*. Whether this is due to different immune responses by hosts or possibly exposure to different lineages of *Bd* is not known [25]. For example, all nine frog species within the *Craugastor punctariolus* clade (robber frogs) [26,27] declined across all their elevational range, from 0 to 2300 meters a.s.l. [28], and yet decades later they appear to be slowly recovering from past *Bd* epizootics [25,29,30]. Similar cases of catastrophic decline followed by apparent recovery have been observed in some highland populations of harlequin frogs, tree frogs and ranid frogs [31,32]. Population fluctuations such as these elicit questions regarding the role of *Bd* transmission, virulence, and lineage in this disease system. Recent studies have shown that *Bd*-GPL is unlikely to be endemic to Costa Rica though it is possible that other endemic *Bd* lineages occur in Costa Rica and throughout the Americas [33].

Retrospective studies analyzing the presence of *Bd* in specimens preserved in natural history collections have been useful to describe *Bd* invasions that may have led to amphibian declines [20,34] as well as situations where *Bd* has been present for a century [10,35,36]. Museum collection data has also contributed to tracking and identifying declined species [37–39]. The utilization and analysis of accurate collections and databases is crucial to understanding the historical context of population declines and can result in more applicable conservation plans. We conducted a retrospective survey using a *Bd* qPCR assay effective on museum specimens [20] to describe the spatial and temporal patterns of *Bd* of anurans in Costa Rica from 1961-2011. We used logistic regression analysis to examine possible environmental factors correlated with *Bd* infection occurrences. Based on our data, we also constructed a habitat suitability model for *Bd* in Costa Rica using a MaxEnt model in order to visualize *Bd* habitat suitability for the region.

## Materials and Methods

### Data collection

We sampled 1016 formalin-fixed, ethanol-preserved museum specimens including thirty-four species of frogs from five taxonomic families. All specimens were collected in Costa Rica between 1961 and 2011 and are housed in the Museum of Zoology (UC Berkeley) and at the Universidad de Costa Rica (UCR). We focused our sampling efforts on anuran species that were reported to have declined during the 1980s and 1990s [40]. Most of the *Craugastor* species have not recovered from population declines and are still classified as critically endangered or extinct according the IUCN [41]. However, we also chose species whose populations initially declined and were subsequently observed to be recovering by around 2010 or later (*Agalychnis annae, Agalychnis lemur, Lithobates vibicarius* and *Lithobates warszewitschii*). The data from our skin swabs, including qPCR results from our survey can be freely accessed on the amphibian disease portal (AmphibiaWeb.org) [42].

### Quantitative PCR Assay

We collected skin swabs from formalin-fixed frogs following a standardized protocol that reduces chances of cross-contamination between specimens [20]. Each museum specimen was removed from the holding jar using flame-sterilized forceps and thoroughly rinsed with 70% ethanol to remove any contaminants from other animals stored in the same jar. Flame-sterilized forceps were used to hold the specimen while swabbing 5 times each of the following locations for a total of 25 strokes, using sterile synthetic cotton swabs; l) the ventral surface from mid abdomen to cloaca, 2) each inner thigh, and 3) the bottom side of the webbing between each toe. Swabs were kept at −4° Celsius until processing in the laboratory. Latex or nitrile gloves were used at all times when handling tubes, jars, and specimens. Gloves were changed between handeling every specimen. *Bd* was extracted from swabs using the Prepman Ultra and Real-Time PCR protocol in the Vredenburg Lab at San Francisco State University [20,43,44]. Positive and negative controls were run in triplicate on every 96-well PCR plate and standard curves were constructed by using 100, 10, 1, and 0.1 *B. dendrobatidis* zoospore quantification standards. Samples were run on an Applied Biosystems 7300 Real-Time PCR thermocycler. We calculated the number of zoospores in terms of Zswab (i.e., estimated *Bd* zoospore genomic equivalents on each swab) by multiplying qPCR results by 80 to account for sample dilution (40 μL Prepman × 10 dilution/ 5 μL for reaction = 80). A *Bd*-positive sample was described as having a Zswab score greater than zero.

### Statistical Analyses

We performed all statistical analyses using the software R (version 3.4.2). To characterize the temporal and spatial occurrence of *Bd* in Costa Rica, we calculated 95% confidence intervals for *Bd* prevalence for each decade sampled based on a binomial probability distribution. We also performed a linear regression for *Bd* infection status as a response variable, assuming a binomial distribution as individuals are either infected or non-infected. Lastly, we used MaxEnt to estimate *Bd* habitat suitability using the significant variables from the linear regression model [45].

For the linear regression, we used the elevation and 19 bioclim variables available on WorldClim (http://www.worldclim.org) and reduced the number of variables by performing a Pearson-correlation test to eliminate highly correlated factors (>0.9 or <-0.9). The following variables were then used; annual mean temperature, mean diurnal temperature range, day-to-night temperature oscillations relative to the annual oscillations (isothermality), temperature seasonality, annual precipitation, precipitation of the wettest month, precipitation of the warmest quarter, precipitation of the driest quarter, and precipitation of the coldest quarter. We then performed a stepwise regression to choose the best-fit model based on the AIC [46,47].

## Results

Our qPCR analysis of the 1016 museum specimens analyzed revealed sixty-eight *Bd*-positive anuran samples and 948 *Bd*-negative anurans for an overall infection prevalence of 6.7% (supplementary table 1). The earliest records were detected in four *Lithobates vibicarius* specimens collected in 1964 from the central volcanic mountain range, on the hillsides of Poas Volcano (fig 1a, supplementary table 2). UCR museum records of the species included in this study begin in the 1960s, thus our retrospective survey begins with the oldest specimens collected in 1961 (n= 2). When grouped by decades we find relatively low *Bd* prevalence in the 1960s and 1970s (4.31%, n= 348 and 4.57%, n= 372, respectively), followed by an increase in the 1980s to 11.70%, n= 171. By the late ’80s and ’90s, *Bd* was detected in museum specimens collected throughout the entire country, whereas earlier positive samples were obtained only from frogs collected in the central regions of the country. Thus, *Bd* became more common throughout the country in the late 1980s and 90s (figure 1a, figure 2). Overall, the majority of the *Bd* positive samples were found in mountains throughout Costa Rica at elevations ranging from 32–2550 meters a.s.l. We found that *Bd* prevalence between species ranged from 0.0% to 46.7%. The species with the highest percentage of *Bd* positive samples came from species that are highly dependent on water for reproduction or live in close proximity to water. For example, we found 46.7% *Bd* infection prevalence (n= 15) in the stream-breeding frog *Hyloscirtus palmeri*, and 45.9% prevalence (n= 37) in *Lithobates vibicarius,* a highland pond-breeding frog. The lowest *Bd* prevalence occurred in the Dendrobatidae and Bufonidae families, in species that spend much of their time on land rather than in water. However, we sampled only a small number of Dendrobatidae specimens (n= 7), and no samples were *Bd*-positive, whereas in Bufonidae we sampled 171 specimens and found that 0.6% were *Bd* positive (supplementary table 1). Overall, the Ranidae family showed the highest percentage of positives (22.4%), followed by Craugastoridae (7.4%). Most Craugastoridae samples were taken from direct developing streamside-breeding species of the *Craugastor punctariolus* clade, which are critically endangered across their entire distribution. *Craugastor andi* however, a species not categorized within the *punctariolus* clade, is also found near streams [28]. In the Craugastoridae family, only four *C. escoces* individuals tested positive out of sixty-three specimens. All *C. escoces* samples were collected before the 1987 Costa Rican amphibian population decline epidemic [24]. The earliest *Bd*-positive *C. escoces* specimens were collected in 1975, and the last was collected in 1986.

**Figure 1a).**
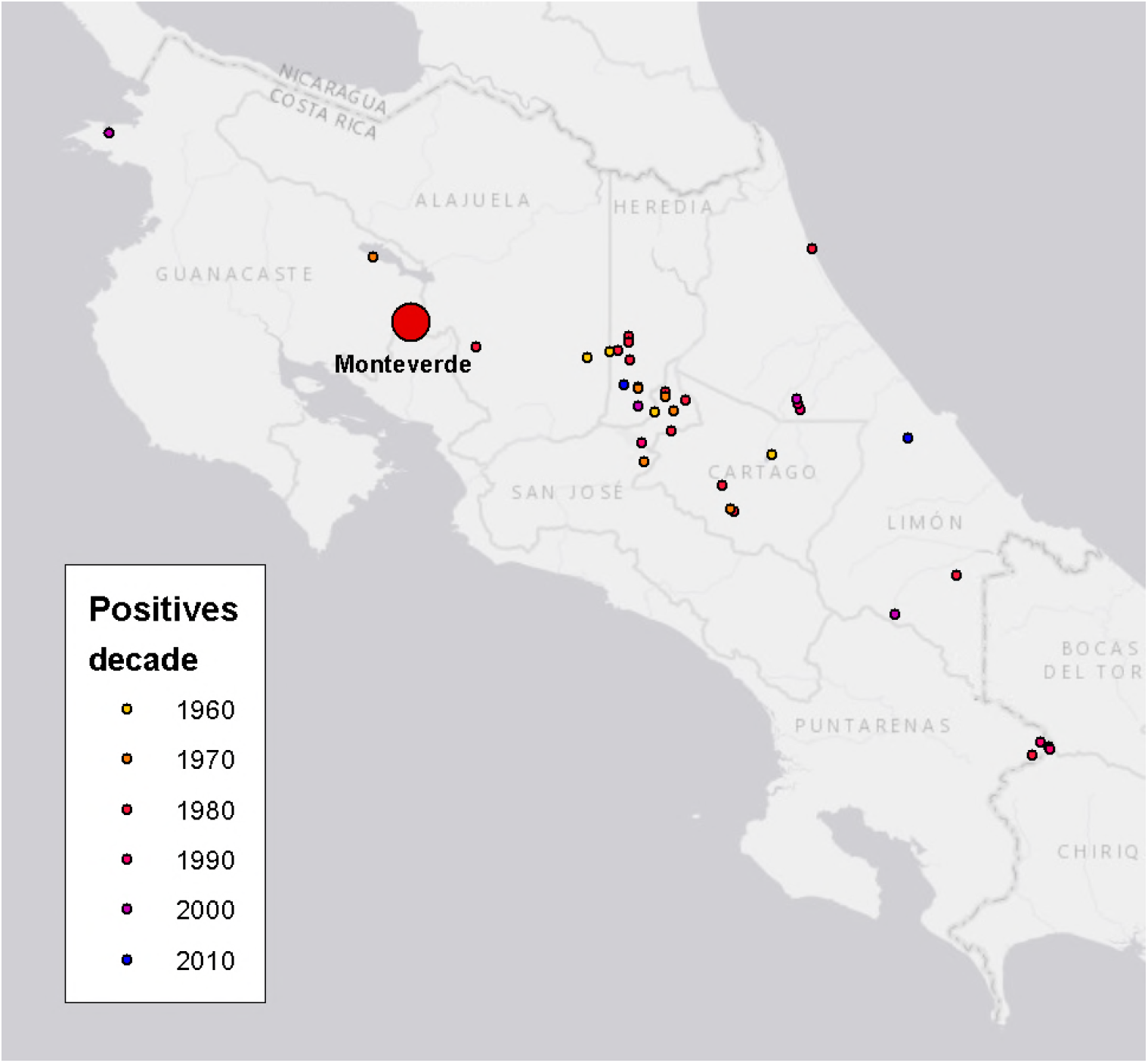
Amphibian museum specimens (1964-2011) that tested positive for *Bd* infection in Costa Rica. Monteverde (red circle) experienced major population declines 1986-89. *Bd*-positive amphibians were collected (n= 32) in the two previous decades in the northern and central mountain range South of Monteverde, but no population studies are available in those areas.

**Figure 1b.**
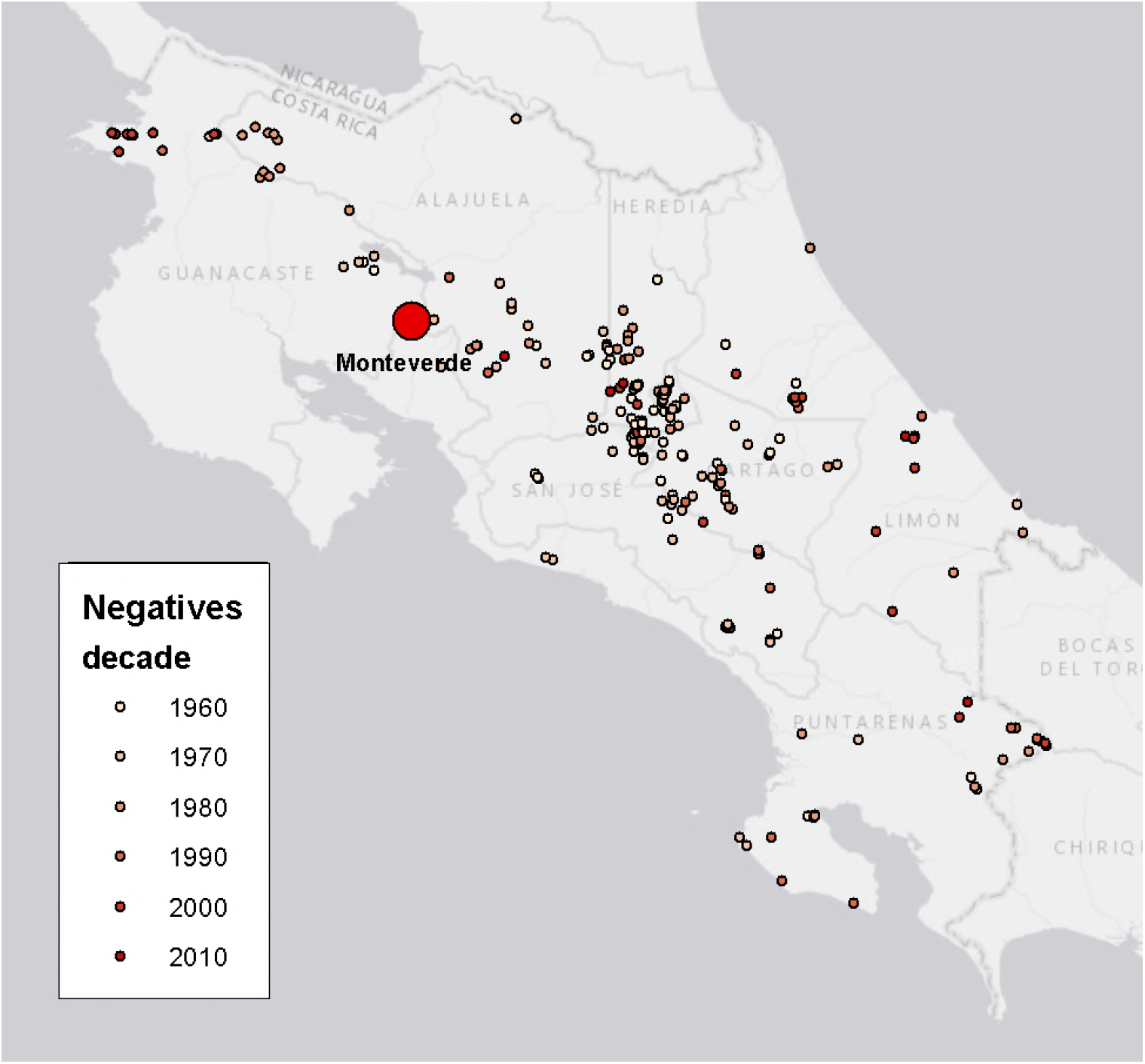
Amphibian museum specimens (1961-2011) that tested negative for *Bd* infection in Costa Rica (n= 948).

**Fig 2.**
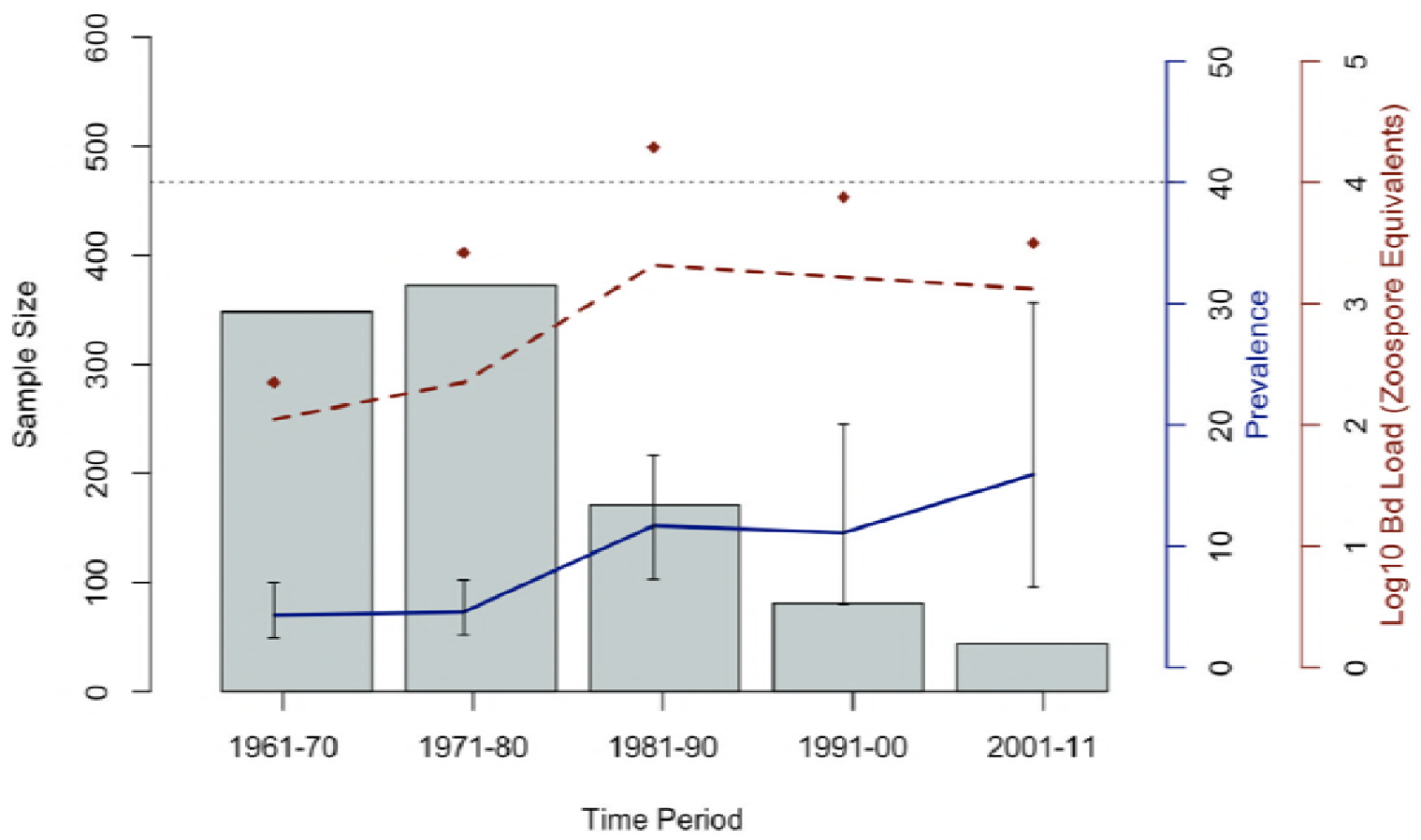
Emergence of *Bd* in Costa Rica from 1961-2011. Infection prevalence (solid blue line) and infection intensity (broken red line) patterns over time along with the maximum zoospore equivalents value per time period (red diamonds). Gray bars represent number of samples analyzed per time period. Dashed line at Log10 Zoospore equivalents = 4 marks the Vredenburg 10,000 value [13].

Our power analysis showed that we had enough samples within each time period to have a robust statistical test (p < 0.01 across all time periods sampled; table 1). In the model with the best AIC (AIC = −2853.44), we found that infection status has a positive relationship with elevation and annual mean temperature (p<0.001 and p<0.001; respectively). We also found that *Bd* infection status has a negative relationship with mean diurnal temperature range (p<0.001). Infection status was not shown to have a significant relationship with the following factors: isothermality, precipitation of the warmest quarter, precipitation of the coldest quarter, precipitation of the wettest month, annual precipitation, temperature seasonality, and precipitation of the driest quarter (table 2). Areas predicted by the *Bd* habitat suitability model to be suitable for *Bd* occurrence, includes mid-elevation ranges across central Costa Rica (fig 3). The areas predicted to be unsuitable include the lowland regions and the coasts.

**Figure 3.**
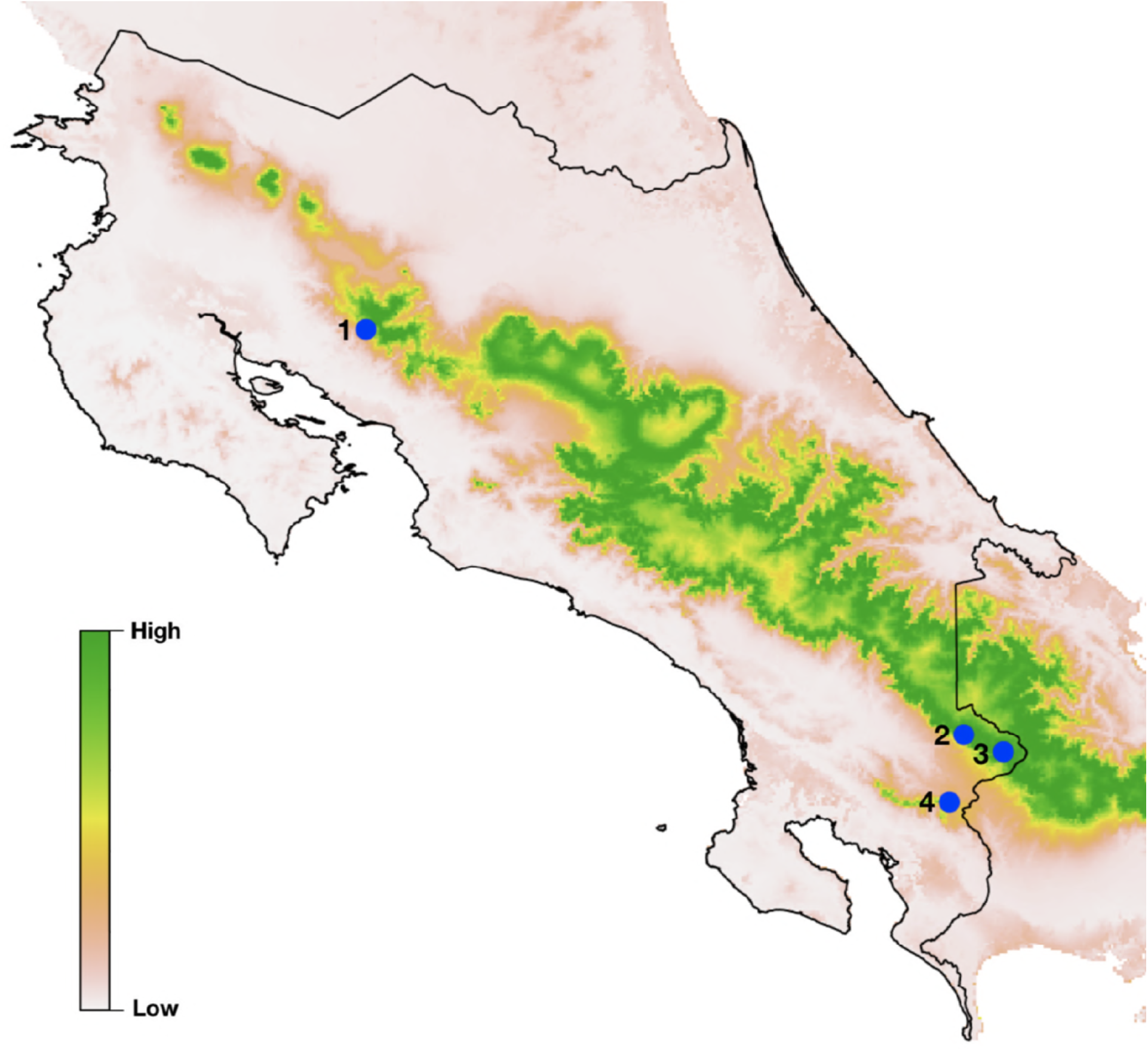
*Bd* habitat suitability model. Areas in Costa Rica predicted to have *Bd* suitability. Increased intensity from white to green indicates increased suitability while blue dots are *Bd*-epizootic localities. (1) Monteverde, where the 1987 decline occurred and (2-4) indicates declines between 1993-1994.

## Discussion

Chytridiomycosis has severely impacted anuran biodiversity worldwide, with hundreds of species affected [3,48]. Though our knowledge of the pathogen is incomplete, our understanding continues to grow through studies of *Bd* pathogen and host dynamics. Our retrospective study revealed that *Bd* was present in Costa Rica twenty-three years before declines were discovered at Monteverde [19]. We found an increasing prevalence of *Bd*-infected hosts collected throughout Costa Rica beginning in the 1970s (fig 1a), with prevalence escalating during and after the known epizootic time period (1980s), which supports the pattern expected with an invasive pathogen.

Previous studies describe the epizootic as a wave that progressed from north-west to south-east Costa Rica and into Panama [20,49,50]. Here, we show a potentially contrasting spatial pattern where *Bd* appears to be present at low prevalence across some of the region (fig 1a) before the purported wave passed through the country. This could be indicative of an invasive pathogen that invades unsuccessfully for decades before epizootics develop, or it could be that the earlier *Bd* infections were the result of a non-virulent lineage of *Bd*, such as has been identified in other parts of the world [35,51,52]. Consistent with an invading pathogen, our limited data show a pattern of spread across the entire time of the study. We found *Bd*-positive specimens only in and around central Costa Rica in the earlier time period (1960s), but by the 1970s, we found *Bd*-positive individuals across a larger area, from north western areas to southern areas of Costa Rica. Our results from the 1980s show the largest expansion of *Bd*, with *Bd*-positive individuals found on the eastern coast of Costa Rica and near the Panama-Costa Rica border to the south-east and all the way north close to Nicaragua. This pattern might reflect the *Bd*-epizootics that are proposed from that time period. In the more recent decades (e.g. 1990s), there were fewer specimens in museum collections that we could test. Thus, constructing a robust statement regarding the spatial distribution of *Bd* in the more recent time period is not possible given the available specimens. Our samples are not free from sampling biases, since museum specimens were collected for reasons unrelated to our study.

In some areas (e.g. South Korea, Brazil, and South Africa), where both the Global Panzootic Lineage (GPL) and an endemic strain of *Bd* occur sympatrically, direct competition between pathogen strains and potential cross immunity of hosts may explain the lack of epizootics [11,53]. Our study provides evidence that *Bd* was present in Costa Rica before the 1987 epizootic in Monteverde, but we acknowledge that there may have been previous undetected epizootics especially since the pathogen was yet described. The few specimens collected in the 1970s showed relatively high levels of infection and, by the 1980s, both prevalence and zoospore equivalents increased (fig 1a, 2). Our data do not refute the studies that show that epizootics in Central America are associated with *Bd* invasion, but they do help provide further data for interpretation. For example, identifying the *Bd* lineage of our earlier positive *Bd* samples would be extremely helpful, since having multiple pathogens that are closely related to each other in a population of hosts may help us understand why *Bd* has had such variable effects on hosts, even in known-epizootic areas. Sub-lethal effects from fighting off the infection of one pathogen can suppress host immunity against other stressors, causing a larger effect [54–56]. However, populations can also benefit by being exposed to a lower virulence pathogen before being exposed to a similar yet more virulent pathogen. Direct competition between pathogens and/or cross immunity has been shown to assist the hosts in acquiring partial or total immunity to one pathogen from a previous infection by another closely related pathogen [57]. Additionally, climate change and the stress of an inconsistent environment may negatively affect amphibians and result in suppressed immune systems, which could make amphibians more vulnerable to chytridiomycosis [69–71]. Future studies involving *Bd* genotyping are required to determine whether *Bd* found in Costa Rica are of a single lineage and whether or not the *Bd* found in epizootics and enzootics are of the same lineage.

Consistent with other studies, our linear regression results found that *Bd* infection status has a positive relationship with elevation and mean temperature [58,59], but contrary to other studies, our best linear regression model (table 2), did not show a relationship between precipitation and *Bd* occurrence [60–63]. This may be due to unintended sampling bias. For example, frogs in the genus *Craugastor* made up a large proportion of available specimens (434 of 1016) and yet most were negative. This genus of frogs are direct developers that do not have a free-swimming aquatic tadpole phase of development, but instead hatch from terrestrial eggs as fully metamorphosed froglets. This more terrestrial lifestyle may decrease exposure to the aquatic pathogen *Bd*, although terrestrial life history alone is not associated with susceptibility to infection [64,65].

Our data show higher *Bd* prevalence in mid to high elevation species, which is possibly due to a more suitable climate for *Bd* [66–68]. The *Bd* habitat suitability model we produced from our data alone, predicts suitable habitat similarly to previous studies [72,73] (fig 3), where the model indicates high habitat suitability for *Bd* where epizootics occurred. For example, in figure 3, we identify the locations with documented *Bd* epizootics occurred in Costa Rica in 1987, 1993, and 1994 (fig 3, blue circles 1-4). Our model also identifies high elevations along the central mountain range as having the highest *Bd* suitability and should be prioritized for further research and monitoring.

The zoospore equivalents (i.e. the Zswab, infection intensity or host infection load) and prevalence of *Bd* observed in this study are typically consistent with epizootic or enzootic dynamics [74]. Nonetheless, our results challenge the hypothesis that *Bd* invaded Costa Rica immediately before epizootics began and suggests that more research is needed to understand and document the role of past potentially failed invasions and/or potential endemic lineages in describing current dynamics of *Bd* and amphibian hosts. Our results show that although *Bd* was present in the 1960s, the significant increase in *Bd*-positive individuals did not begin in the samples available until the 1980s (when the epizootics began). The data we provide in this study are not well-suited to test the novel vs endemic pathogen hypotheses for *Bd* (fig 3) [17]; however, these samples could be used to test for *Bd* lineage in future studies that may shed light on this question [51]. The steady increase in prevalence of *Bd* throughout all elevations in Costa Rica after 1990 suggests that *Bd* has become more broadly established throughout the country [25,75] than it was previously. The recent rediscovery of some remnant populations of frogs once thought extinct provides new opportunities to assess the current impact of *Bd* in highly susceptible species [25,29,30,32]. Consistent with previous studies, we propose that *Bd* epizootics in amphibians began in the central range of Costa Rica, affecting stream-breeding and pond-breeding species that inhabited this region (supplementary tables 1 and 2) such as *Lithobates vibicarius, Isthmohyla angustilineata, I. tica, I. xanthosticta, I. rivularis, Duellmanohyla uranochroa Craugastor fleischmanni, C. ranoides, C. escoces, C. sp.* (*C. punctariolus* clade), *C. melanostictus, C. andi, Atelopus varius, A. senex* (Harlequin frogs), and *Incilus holdrigei*.

In this study we discovered *Bd*-infected frogs in Costa Rica twenty-three years before enigmatic amphibian declines occurred. These infected animals could represent failed pathogen invasions (e.g. “pathogen fade out” theory [76,77]), slower than expected invasion dymaics resulting in epizootics, or endemic lineages of *Bd* that may exhibit enzootic pathogen/host dynamics. New studies that sequence and identify *Bd* lineages from our data could help create a more complete understanding of the lineage and spread patterns of *Batrachochytrium dendrobatidis* in Central America.

## Acknowledgements

We thank the Museum of Vertebrate Zoology for access to their natural history collections. We are very grateful to the numerous undergraduate and graduate student volunteers who assisted in lab work at the Vredenburg Lab at San Francisco State University.

---

**Table 1. Batrachochytrium dendrobatidis (***Bd***) prevalence in museum specimens collected in Costa Rica**. Pr (no *Bd*) is the probability of finding no *Bd*-positive samples in each time period if *Bd* was present with an enzootic prevalence of 11.0% [35].

**Table 2. Linear regression output. Environmental factors and their relationship to *Bd* infection status**. (+) next to the variable name indicates a positive relationship between factor and *Bd* presence. (-) indicates a negative relationship between factor and *Bd* presence. The model with the lowest AIC value (1st model; AIC = −2853.44) was considered the best model.

**Supplementary Table 1.**
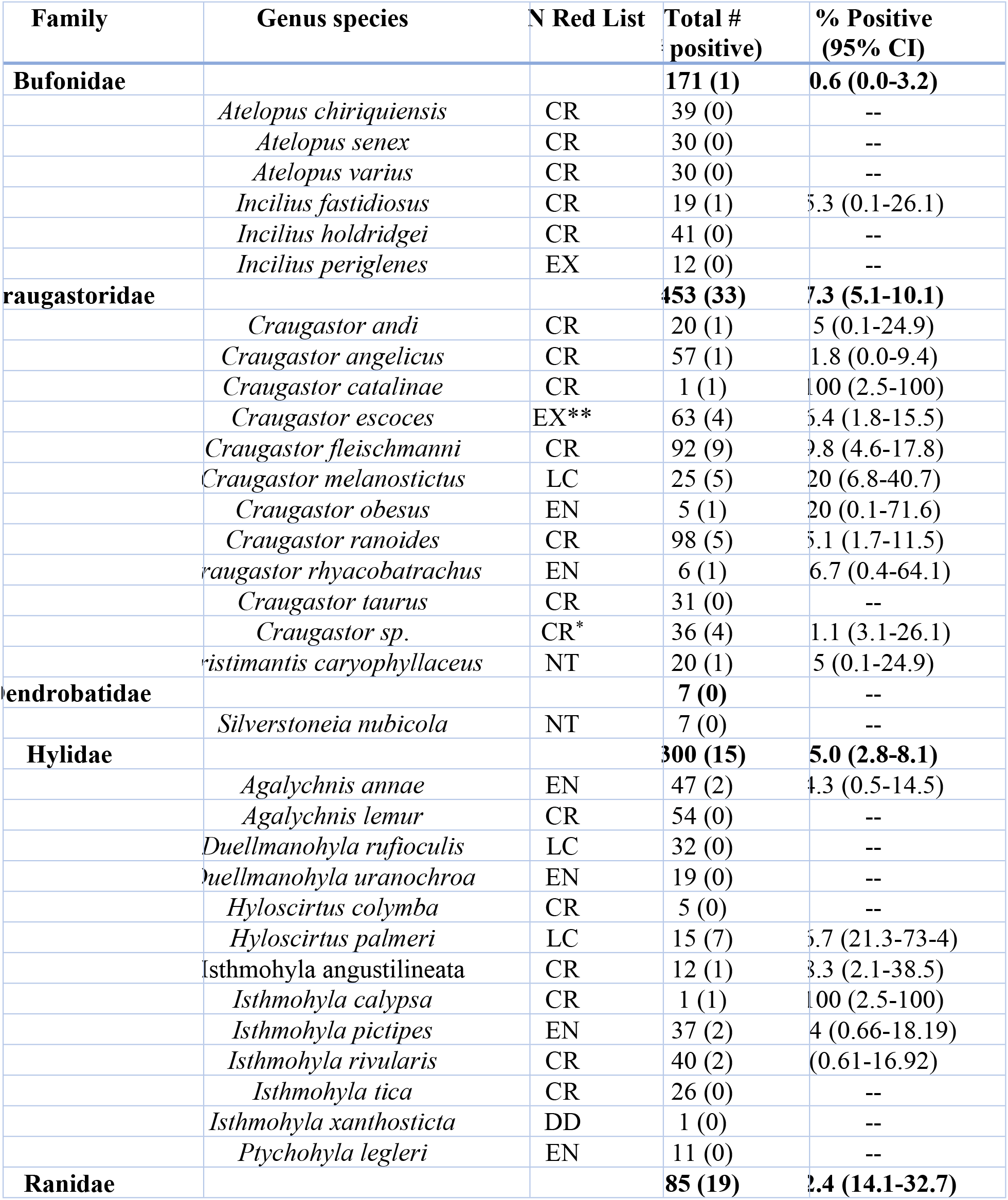
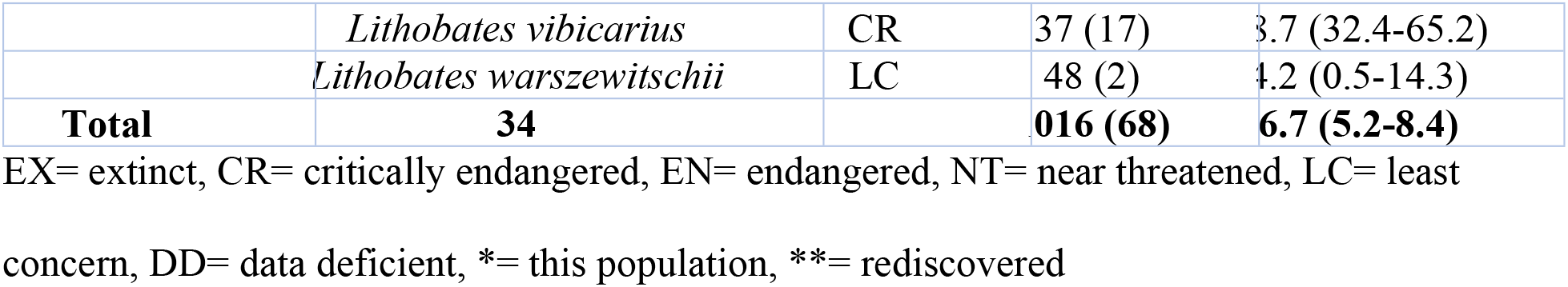
*Bd* observed in in museum specimens from Costa Rica. The table shows surveyed species, conservation status and proportion of samples with *Bd* including 95% binomial confidence intervals.

**Supplementary Table 2.** Geographic and time data for all *Bd*-positive species surveyed in UCR museum specimens from Costa Rica.

| Species | # Positive | Year collected | Elevation (m) | Localities<sup>+</sup> |
| --- | --- | --- | --- | --- |
| *Incilius fastidiosus* | 1 | 1990 | 2400 | Las Tablas |
| *Craugastor andi* | 1 | 1986 | 1300 | La Virgen |
| *Craugastor angelicus* | 1 | 1967 | 1370 | Cinchona |
| *Craugastor catalinae* | 1 | 1990 | 1800 | Las Tablas |
| *Craugastor escoces* | 4 | 1975(3), 1986 | 1500, 2000 (3) | Chompipe (3), PNBC |
| *Craugastor fleischmanni* | 9 | 1974(2), 1975(2), 1977, 1982, 1984, 1986, 2010 | 900, 1260, 1300, 1700, 1800, 2000, 2011, 2110 (2) | Chompipe (3), Desamparados, Chompipe, Honduras, San Ramón (2), Porrosati |
| *Craugastor melanostictus* | 5 | 1968 (2), 1970 (2), 1986 | 1500, 2000 (4) | Chompipe (4), Varablanca |
| *Craugastor obesus* | 1 | 1984 | 700 | Nimaso |
| *Craugastor ranoides* | 5 | 1968, 1985 (2), 1986, 2009 | 1 (2), 32, 600, 900 | Tortuguero (2), Reserva, Murciélago, Turrialba |
| *Craugastor rhyacobatrachus* | 1 | 1983 | 1260 | Mellizas |
| *Craugastor sp.* | 4 | 1971, 1984 (3) | 1240, 1600, 1640 | Tapantí |
| *Pristimantis caryophyllaceus* | 1 | 2007 | 1500 | Telire |
| *Agalychnis annae* | 2 | 1992, 2004 | 1150, 1500 | San Francisco, Concepción |
| *Hyloscirtus palmeri* | 7 | 1984, 1997, 1999 (3), 2002, 2011 | 360, 400, 600, 775 (3), 1400 | Guayacán (5), Veragua, Carrillo |
| *Isthmohyla angustilineata* | 1 | 1984 | 1600 | Tapantí |
| *Isthmohyla calypsa* | 1 | 2002 | 2300 | Las Tablas |
| *Isthmohyla pictipes* | 2 | 1969, 1971 | 2000, 2110 | Chompipe (2) |
| *Isthmohyla rivularis* | 2 | 1984, 1990 | 1240, 1800 | Tapantí, Las Tablas |
| *Lithobates vibicarius* | 17 | 1964 (4), 1966, 1968 (5), 1971 (2), 1976, 1979, 1984, 1986, 1990 | 1400, 1500, 1700, 2000 (5), 2050, 2110 (3), 2400, 2550 (4) | Poás (4), Chompipe (8), San Jerónimo, Cascajal (2), Varablanca, Las Tablas |
| *Lithobates warszewitschii* | 2 | 1970, 1984 | 650, 1400 | Tilarán, La Mula |

